# Engineering transcription factor BmoR mutants for constructing multifunctional alcohol biosensors

**DOI:** 10.1101/2021.10.21.465235

**Authors:** Tong Wu, Zhenya Chen, Shuyuan Guo, Cuiying Zhang, Yi-Xin Huo

## Abstract

Native transcription factor-based biosensors (TFBs) have the potential for in situ detection of value-added chemicals or byproducts. However, their industrial application is limited by their ligand promiscuity, low sensitivity, and narrow detection range. Alcohols exhibit similar structures, and no reported TFB can distinguish a specific alcohol from its analogs. Here, we engineered an alcohol-regulated transcription factor, BmoR, and obtained various mutants with remarkable properties. For example, the generated signal-molecule-specific BmoRs could distinguish the constitutional isomers n-butanol and isobutanol, with insensitivity up to an ethanol concentration of 800 mM (36.9 g/L). Linear detection of 0–60 mM of a specific higher alcohol could be achieved in the presence of up to 500 mM (23.0 g/L) ethanol as background noise. Notably, two mutants with raised outputs and over 1.0 × 10^7^-fold higher sensitivity, and one mutant with an increased upper detection limit (14.8 g/L n-butanol or isobutanol) were screened out. Using BmoR as an example, this study systematically explored the ultimate detection limit of a TFB towards its small-molecule ligands, paving the way for in situ detection in the biofuel and wine industries.

## Introduction

Upon sensing specific signal molecules, transcription factors (TFs) can bind or unbind to the DNA-regulatory sequences of target genes to induce or repress gene transcription. TFs can be utilized as biosensors by coupling the transcriptional alteration with the expression of a reporter protein^1^. Transcription factor-based biosensors (TFB) have been designed and constructed to detect toxic metals ^2^, regulate metabolic flux ^3, 4^, and screen highly active enzymes ^5^ or chemical overproducers ^6, 7^. However, several drawbacks limit the industrial applications of TFBs. First, TFB signaling can be saturated by the medium concentration of intracellular signal products and cannot distinguish chemical overproducers from producers ^8^. Second, TFBs can be interfered by the byproducts, which are structurally similar to the products ^9^. Third, in many cases, TFBs are not sensitive enough to detect low concentrations of value-added chemicals^10, 11^. Therefore, it is necessary to develop TFBs with low ligand promiscuity, high sensitivity, and a wide detection range.

n-Butanol and isobutanol are considered as promising substitutes for gasoline because of their special properties^12^. Butanols are also crucial precursors for the production of plastics and polymers^13^. Metabolic engineering has been used to produce n-butanol and isobutanol in various hosts ^14–22^. However, an efficient TFB with low ligand promiscuity, high sensitivity, and a wide detection range has yet to be achieved for in situ detection and high-throughput screening. *Pseudomonas butanovora* BmoR is an activated transcription factor that belongs to the bacterial enhancer-binding proteins (bEBPs) and can induce the transcription of σ^54^-dependent *P_bmo_* promoter by combining with C2-C5 n-alcohols ^23, 24^. The activation mechanism of BmoR is shown in Fig. 1A. The N-terminal, central domain, and C-terminal of BmoR are responsible for sensing and combining the signal molecules, contacting the σ^54^ factor of holoenzyme and hydrolyzing ATP, and binding to the upstream sequence of the σ^54^-dependent promoter, respectively ^24^. Native BmoR has been shown to respond to C3-C5 alcohols, and a BmoR-based biosensor has been utilized to screen n-butanol or isobutanol producers ^10^.

**Fig. 1.**
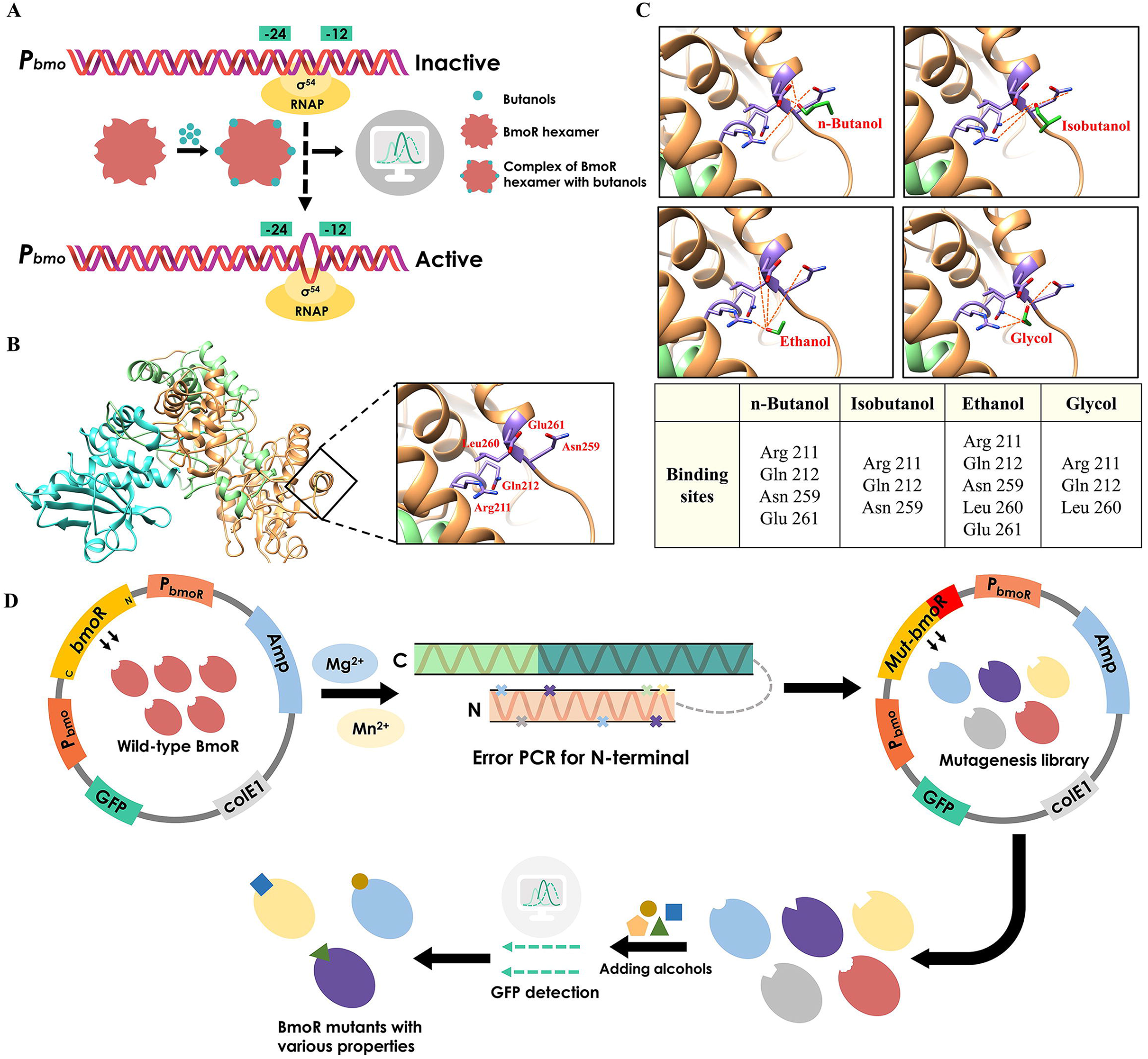
The activation mechanism and simulated structure of wild-type BmoR. **(A)** The simulation mechanism of wild-type BmoR. **(B)** The simulated structure and the binding pocket of wild-type BmoR. **(C)** The binding pocket and binding sites of wild-type BmoR with different signal molecules (isobutanol, n-butanol, ethanol or glycol). **(D)** The whole screening process of the N-terminal mutagenesis library.

In this study, we aimed to obtain efficient BmoR mutants with specificity, high sensitivity, and a wide detection range towards special higher alcohols for the purpose of in situ detection and high-throughput screening. The development of a biosensor that was sensitive to butanols but completely insensitive to 0–36 g/L ethanol was another significant purpose of this study. This type of biosensor is urgently needed in the wine industry for the in situ detection of byproducts (n-butanol or isobutanol) production without interfering with ethanol production. Here, we utilized error-prone PCR to construct a BmoR random mutagenesis library based on the understanding of the BmoR N-terminal and a variety of BmoR mutants with desired properties, such as specificity, sensitivity or a wide detection range were obtained and are shown in Table 1. These mutants could be widely utilized in various fields, including metabolic regulation, enzyme or production strain screening, and byproduct detection.

**Table 1.** Representative BmoR mutants with remarkable properties in this study

| BmoR<br>Mutant | Property |  |  |  |  |  |
| --- | --- | --- | --- | --- | --- | --- |
|  | n-Butanol<br>specificity | Isobutanol<br>specificity | Insensitivity<br>to ethanol | Higher<br>sensitivity | Wider<br>detection<br>range | Higher<br>output |
| W21R/E54V | √ |  | √ |  |  |  |
| I183T/D273N | √ |  | √ |  |  |  |
| M94V/F272L |  | √ | √ |  |  |  |
| S240P |  | √ | √ |  |  |  |
| E54G |  |  |  | √ |  | √ |
| V311A |  |  |  | √ |  | √ |
| T12N |  |  |  |  | √ |  |

## Results and discussion

### Analysis of the wild-type BmoR structure and establishment of the mutagenesis library

We first analyzed the binding regions of different molecules and attempted to utilize rational design to endow BmoR with signal-molecule specificity. In our previous study, the biofilm regulator FleQ (PDB code: 5EXP) and the transcription factor PobR (PDB code: 5W1E) were used as templates to simulate wild-type BmoR (Fig. 1B). In this study, molecular docking of BmoR with different alcohols (n-butanol, isobutanol, ethanol, and glycol) was conducted. The results in Fig. 1C show that the binding pockets of different signal molecules were similar, and the signal molecules bound to BmoR mainly via the formation of hydrogen bonds with Arg211, Gln212, and Asn259. We first performed site-saturation mutagenesis at these sites. The generated mutants had decreased response values when compared to the wild-type (Fig. S1A, B, and C) and did not display signal-molecule specificity. These results suggest the complexity of the response mechanisms of BmoR and indicate that obtaining BmoR with desirable properties cannot be easily achieved through rational design. Therefore, we performed random mutagenesis of BmoR. The N-terminus of BmoR is responsible for recognizing and binding signal molecules and modulating the activity of bEBP ^25, 26^. Modifying the region could directly influence the response of BmoR to the signal molecule and further change its properties.

Therefore, error-prone PCR was conducted at the N-terminal of BmoR to endow BmoR with desirable properties. The entire screening process is illustrated in Fig. 1D. After screening the mutagenesis library, we acquired 400 colonies that had different GFP values when compared with wild-type BmoR. The *bmoR* genes in these colonies were then extracted and sequenced.

Besides, we assumed the initiation mechanism of BmoR. BmoR and NtrC could both activate the σ^54^-dependent transcription. The regulatory mechanism of NtrC was positive regulation^27^. In our previous study we have proved that the regulatory mechanism of BmoR was also positive regulation^24^. Therefore, we speculated the initiation mechanism of BmoR might be similar to NtrC because of their structural and functional similarity. To induce the transcription, BmoR might be firstly expressed and formed as dimers. The dimers of BmoR would then capture the signal molecules and the σ^54^ of σ^54^-RNAP holoenzyme (Eσ^54^) would bind to the −24 (GG) and −12 (TGC) regions of σ^54^-dependent promoter (*P_bmo_*). After that, the dimers with signal molecules would form as hexamer to initiate the transcription of *P_bmo_* with the energy which would be released by ATP hydrolysis. The whole activation process was shown in Fig. S2. The true regulation mechanism of BmoR needed to be exploited by crystallization of BmoR with signal molecules and in-depth analysis of the generated structures.

### Specific response of BmoR mutants towards n-butanol

We isolated two special mutants (W21R/E54V and I183T/D273N) that show a significant response towards 10 mM of n-butanol and a weak response towards 10 mM of isobutanol (Fig. 2A). The GFP/OD_600_ values of W21R/E54V and I183T/D273N towards 10 mM of n-butanol were 92.4 ± 14 and 15.6 ± 0.8, respectively (Fig. 2B). A series of gradient addition experiments with different concentrations of n-butanol or isobutanol were carried out to measure the apparent *K*_m_ value and to verify the detection potential of these two mutants. Significantly, the GFP/OD_600_ values of W21R/E54V and I183T/D273N increased with the increase of n-butanol concentration (Fig. 2C), and the response values of W21R/E54V and I183T/D273N reached 325 ± 17 and 147 ± 12 when 100 mM n-butanol was added to the culture, respectively, while the response values of these two mutants have no obvious change with the increase of isobutanol concentration (Fig. 2D). These results suggest that W21R/E54V and I183T/D273N could perceive the change of n-butanol concentration and further give the corresponding response, and exhibited insensitivity towards isobutanol. In addition, the apparent *K*_m_ value of W21R/E54V was 42.6 mM towards n-butanol. The response value of I183T/D273N was linear to the concentration of n-butanol. Based on the above results, we verified the mutants W21R/E54V and I183T/D273N as n-butanol-specific BmoR mutants.

**Fig. 2.**
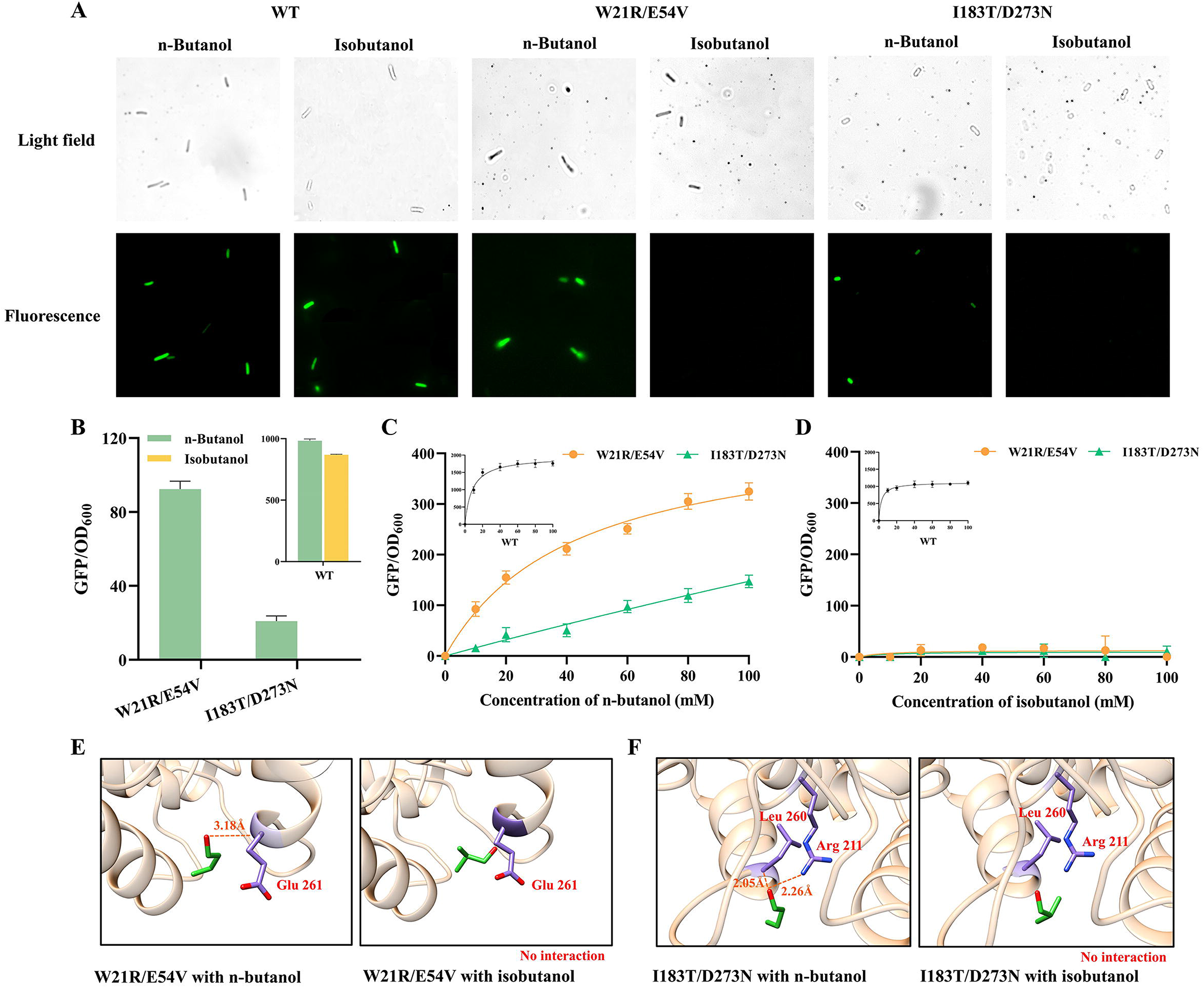
Specific response of BmoR mutants towards n-butanol. **(A)** The green fluorescence of mutants W21R/E54V and I183T/D273N towards 10 mM n-butanol or isobutanol. **(B)** The response values of BmoR mutants W21R/E54V and I183T/D273N towards 10 mM n-butanol or isobutanol. **(C)** The response curves of BmoR mutants towards n-butanol. **(D)** The response curves of BmoR mutants towards isobutanol. **(E)** Simulation and molecule docking of mutant W21R/E54V with n-butanol or isobutanol, and confirmation of the binding sites with signal molecules. **(F)** Simulation and molecule docking of mutant I183T/D273N with n-butanol or isobutanol, and confirmation of the binding sites with signal molecules. Values and error bars represent mean and s.d. (n = 3), respectively.

In addition, simulation and molecular docking of W21R/E54V and I183T/D273N with isobutanol or n-butanol were conducted. The model in Fig. 2E shows that a hydrogen bond was formed between n-butanol and the 261^st^ glutamate of W21R/E54V, and there was no force between isobutanol and W21R/E54V, suggesting that the interaction between n-butanol and W21R/E54V was stronger than that between isobutanol and W21R/E54V. In addition, we mutated the 261^st^ glutamate to the other 19 amino acids. The generated 19 mutants have significantly lower response values when compared with W21R/E54V, which demonstrates the significance of Glu261 in the response to n-butanol (Fig. S3). Similar to W21R/E54V, two hydrogen bonds between n-butanol and I183T/D273N were observed in the complex, and no force was formed between isobutanol and I183T/D273N, demonstrating that n-butanol could bind I183T/D273N tightly (Fig. 2F).

### Specific response of BmoR mutants towards isobutanol

In addition to the n-butanol-specific BmoR mutants, two mutants (M94V/F272L and S240P) that significantly responded to 10 mM isobutanol and hardly responded to 10 mM n-butanol, were screened from the mutagenesis library (Fig. 3A and B). The results of the gradient addition experiment in Fig. 3C and D show that the GFP/OD_600_ values of M94V/F272L and S240P exhibited an increasing trend with the increase of isobutanol concentration and had no significant change with the increase of n-butanol concentration. These distinct responses illustrated that M94V/F272L and S240P have the ability to perceive the change of isobutanol concentration and then output the corresponding response signal; on the contrary, they could not sense n-butanol. Hence, the mutants M94V/F272L and S240P could be considered as isobutanol-specific BmoR mutants.

**Fig. 3.**
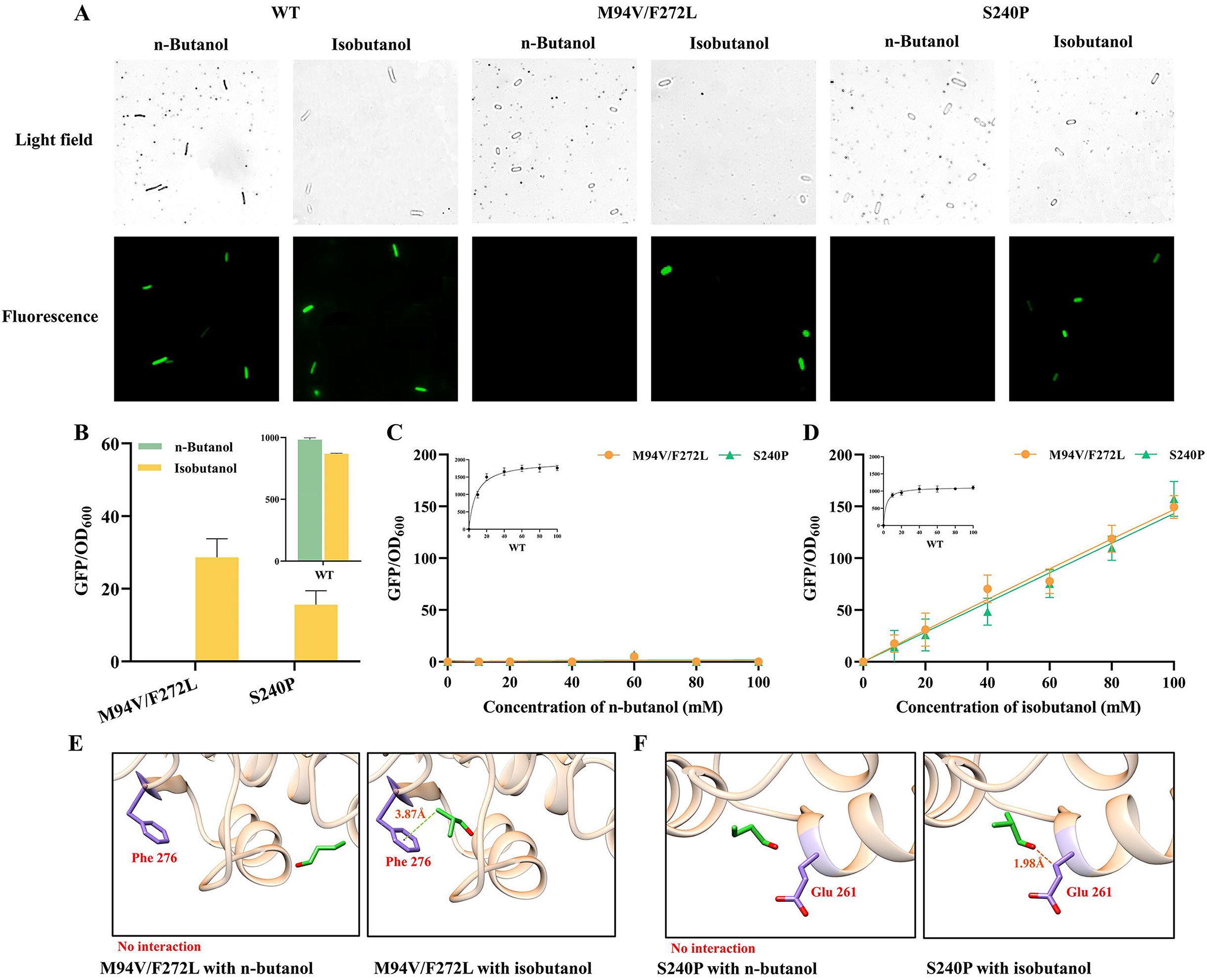
Specific response of BmoR mutants towards isobutanol. **(A)** The green fluorescence of mutants M94V/F272L and S240P towards 10 mM n-butanol or isobutanol. **(B)** The response values of BmoR mutants M94V/F272L and S240P towards 10 mM n-butanol or isobutanol. **(C)** The response curves of BmoR mutants towards n-butanol. **(D)** The response curves of BmoR mutants towards isobutanol. **(E)** Simulation and molecule docking of mutant M94V/F272L with n-butanol or isobutanol, and confirmation of the binding sites with signal molecules. **(F)** Simulation molecule docking of mutant S240P with n-butanol or isobutanol, and confirmation of the binding sites with signal molecules. Values and error bars represent mean and s.d. (n = 3), respectively.

Furthermore, complexes containing any of the above mutants and signal molecules (n-butanol or isobutanol) were modeled, and the interactions between mutated BmoR and the signal molecule were analyzed to explain the isobutanol specificity of M94V/F272L and S240P. As shown in Fig. 3E, hydrophobic interactions existed between the signal molecule isobutanol and Phe276 of M94V/F272L; meanwhile, no interaction was observed in the complex of M94V/F272L and n-butanol. For the complexes of mutant S240P and signal molecules, isobutanol bound to S240P via the formation of a hydrogen bond at Glu261 and n-butanol did not interact with S240P (Fig. 3F). The interaction between the isobutanol-specific mutants and signal molecules proved the isobutanol specificity of the mutants.

### Further verification of the specificity via the addition of mixed signal molecules

To further confirm the specificity and test whether the two kinds of signal molecules competed to combine the mutants, the response of n-butanol-specific and isobutanol-specific BmoR mutants was measured by adding mixed signal molecules to the culture. We used an n-butanol-specific mutant (W21R/E54V) and an isobutanol-specific mutant (M94V/F272L) as examples to conduct the experiments. As shown in Fig. 4A, the GFP/OD_600_ value of the n-butanol-specific mutant W21R/E54V increased with an increase in the n-butanol proportion, reaching 342 ± 7 when the ratio of n-butanol to isobutanol in the culture was 10:1 (100 mM:10 mM). As a comparison, the GFP/OD_600_ value of W21R/E54V did not change with the increase of isobutanol proportion. The response of W21R/E54V only varied with the change of n-butanol proportion, proving that there is no competition between n-butanol and isobutanol in the n-butanol-specific mutant and that W21R/E54V could maintain its specificity in the case of mixed signal molecules. Similar to W21R/E54V, the GFP/OD_600_ value of M94V/F272L only varied with the change of the isobutanol proportion. In summary, we screened n-butanol-specific BmoR mutants (W21R/E54V and I183T/D273N) and isobutanol-specific BmoR mutants (M94V/F272L and S240P) from the mutagenesis library, and these mutants showed satisfactory specificity.

**Fig. 4.**
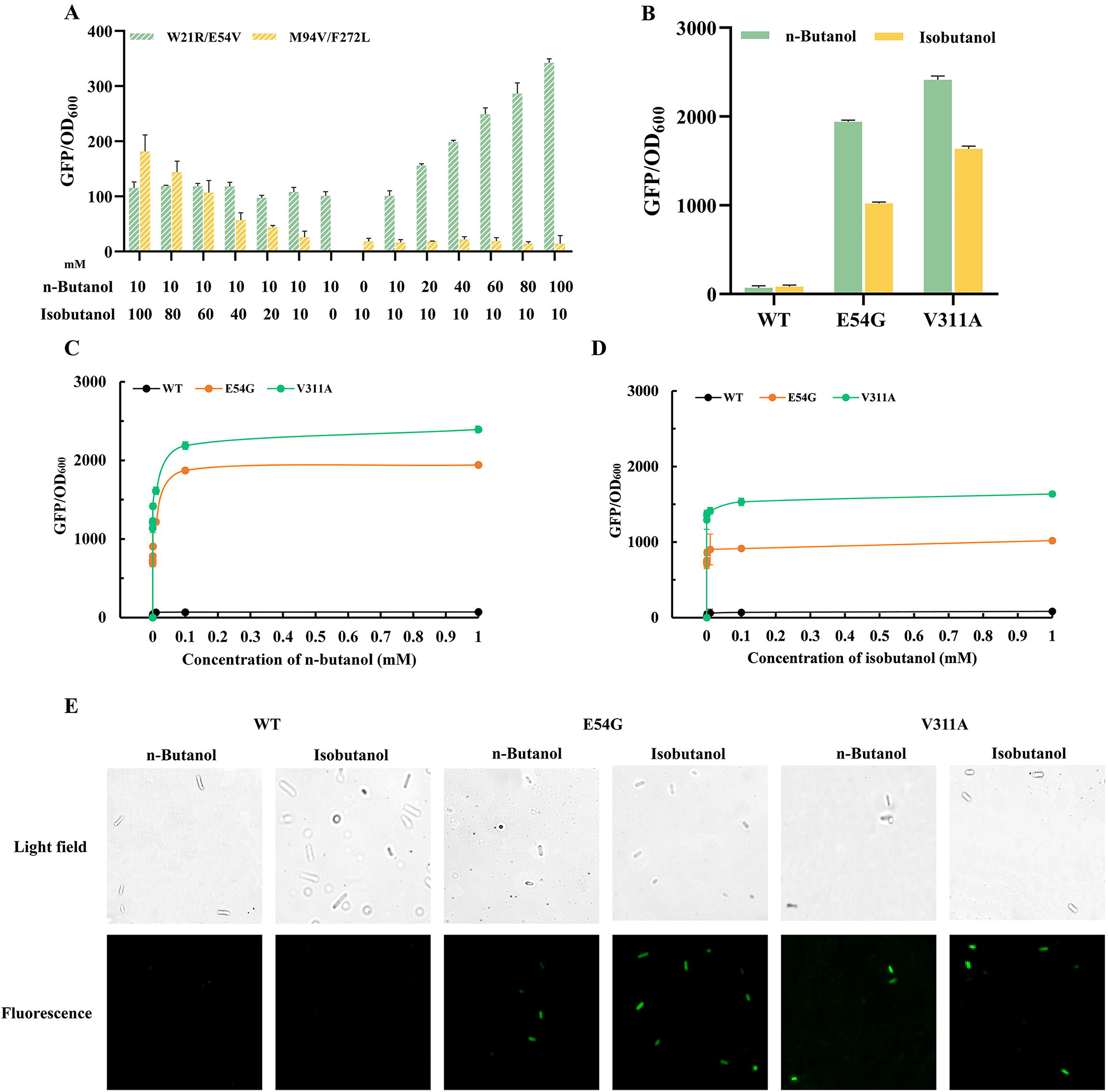
Verification of the specificity via adding mixed signal molecules and BmoR mutants with higher outputs and sensitivity. **(A)** Maintaining the concentration (10 mM) of one kind of butanols and increasing the concentration of the other kind of butanols with a gradient (0-100 mM) to confirm the specificity of BmoR mutants W21R/E54V and M94V/F272L. **(B)** The response values of BmoR mutants E54G and V311A towards 1 mM n-butanol or isobutanol. **(C)** The response curves of BmoR mutants E54G and V311A towards n-butanol. **(D)** The response curves of BmoR mutants E54G and V311A towards isobutanol. **(E)** The green fluorescence of mutants E54G and V311A towards 1 × 10^−6^ mM n-butanol or isobutanol. Values and error bars represent mean and s.d. (n = 3), respectively.

### BmoR mutants with higher output and sensitivity

In addition to the above signal molecule-specific mutants, two mutants with higher outputs were isolated from the random mutagenesis library. The sequencing results show that these two mutants have mutations of E54G and V311A, respectively. Notably, mutant E54G have a 27.5-fold higher GFP/OD_600_ value (1941 ± 17) towards 1 mM n-butanol and a 12.1-fold higher GFP/OD_600_ value (1020 ± 17) towards 1 mM isobutanol when compared with wild-type BmoR. In comparison, the GFP/OD_600_ values of V311A towards 1 mM n-butanol and 1 mM isobutanol were 34.2- and 19.5-fold higher than wild-type BmoR, respectively (Fig. 4B). Based on this, we assumed that these two mutants with higher output might respond to the low concentration of butanols, signifying that the mutants have higher sensitivity. To validate this assumption, we added 0–1 mM n-butanol or isobutanol to the culture to test the response of mutants E54G and V311A. The lower detection limit of BmoR was defined as the concentration of butanols that could achieve a maximum GFP/OD_600_ value of 75%. Based on this, we estimated the lower detection limits of mutants E54G and V311A based on the Michaelis-Menten equation of Origin8.5. As shown in Fig. 4C and D, the lower detection limits of mutants E54G and V311A towards n-butanol were 2.64 × 10^−6^ mM and 2.13 × 10^−6^ mM, respectively, which demonstrated that the sensitivity of these two mutants towards n-butanol was over 1.0 × 10^7^-fold higher than that of wild-type BmoR (28 mM). In comparison, mutants E54G and V311A have lower detection limits of 2.16 × 10^−6^ mM and 1.94 × 10^−6^ mM towards isobutanol, respectively, which suggests that these two mutants possess over 1.0 × 10^7^-fold higher sensitivity to isobutanol when compared with wild-type BmoR (26 mM). After that, we fed 1 × 10^−6^ mM of n-butanol or isobutanol to the cultures of mutants E54G and V311A, and used fluorescence microscope to observe the green fluorescence. As shown in Fig. 4E, mutants E54G and V311A have significant green fluorescence towards 1 × 10^−6^ mM of n-butanol or isobutanol. As a comparison, wild-type BmoR has no obvious green fluorescence. These results suggested mutants E54G and V311A possess high sensitivity to n-butanol and isobutanol. In addition, we attempted to introduce the mutations in the mutant with high output into the signal molecule-specific mutants in order to enhance the response values of the signal molecule-specific mutants. However, the mutations in the mutant with high output did not significantly enhance the output of signal molecule-specific mutants (data not shown). Future work might focus on enhancing the sensitivity and output of signal molecule-specific mutants in order to sense the trace accumulation of butanols.

### BmoR mutant with wider detection range

To utilize the BmoR-based biosensor to screen industrial high-level n-butanol or isobutanol production strains, we needed to broaden the detection range of BmoR. In this study, we screened out a mutant (T12N) with a wider detection range (0–200 mM) when compared with wild-type BmoR. Fig. 5A and B showed mutant T12N outputted significantly distinct GFP/OD_600_ values when feeding the 150 mM and 200 mM butanol solutions and the apparent *K*_m_ values of T12N were 37.3 mM towards n-butanol and 89.9 mM towards isobutanol. These results suggest that the upper detection limit of mutant T12N could reach 200 mM.

**Fig. 5.**
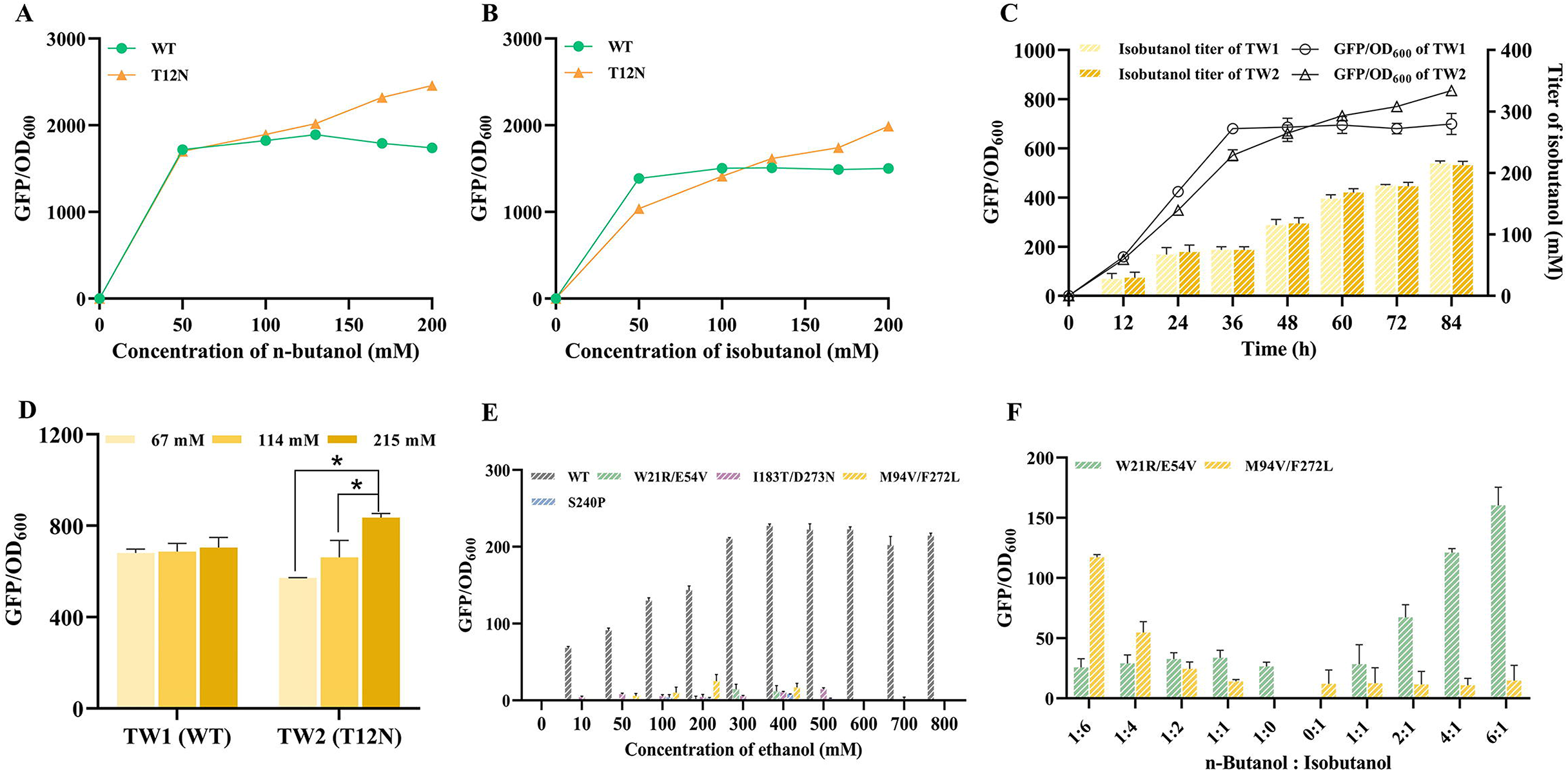
BmoR mutant with wider detection range or ethanol insensitivity. **(A)** The response curves of wild-type BmoR and mutant T12N towards n-butanol. **(B)** The response curves of wild-type BmoR and mutant T12N towards isobutanol. **(C)** The response curves of wild-type BmoR and mutant T12N towards isobutanol which was produced by the isobutanol-producing strain. **(D)** The response values of wild-type BmoR and mutant T12N towards 67 mM, 114 mM and 215 mM isobutanol. **(E)** The response values of specific mutants via adding 0-800 mM ethanol in culture. **(F)** The response values of W21R/E54V and M94V/F272L via adding 0-60 mM n-butanol or isobutanol and 500 mM ethanol as noise. Values and error bars represent mean and s.d. (n = 3), respectively. *P < 0.1, **P < 0.01 as determined by two-tailed t-test.

We then introduced mutant T12N into an isobutanol-producing strain to confirm its wider detection range for *in vivo* host-producing isobutanol. First, plasmids pYH10 and pYH10-T12N were individually introduced into the isobutanol-producing strain TW, resulting in strains TW1 and TW2, respectively. TW1 and TW2 were used for subsequent fermentation and GFP/OD_600_ value detection. As shown in Fig. 5C and D, the GFP/OD_600_ value of strain TW2 with T12N BmoR expression displayed an upward trend as the isobutanol titer during the 84-hour fermentation. The GFP/OD_600_ values at 72 h and 84 h were significantly different, and the value at 84 h reached 835 ± 25. Meanwhile, 215 mM isobutanol was accumulated in the culture. In comparison, the GFP/OD_600_ value of strain TW1 with wild-type BmoR expression increased with the production of isobutanol in the first 36 h, and the isobutanol titer was 67 mM at 36 h. In the next 48 h, isobutanol was continuously produced. However, the GFP/OD_600_ value of TW1 did not increase significantly. These results illustrate that mutant T12N demonstrates a wider detection range (0–200 mM) towards *in vivo* host-producing isobutanol, and this mutant could be used to construct a sensor to screen overproducers.

### BmoR mutants with ethanol insensitivity

In the ethanol fermentation industry, n-butanol and isobutanol are usually produced as byproducts. We assumed that the BmoR-based biosensor could serve as a powerful tool to detect the accumulation of higher alcohols in ethanol-producing strains. However, native BmoR showed a significant response to ethanol, which could induce signal interference and lead to inaccurate measurement of higher alcohol concentrations. Therefore, a biosensor that is insensitive to ethanol and sensitive to higher alcohols needs to be urgently explored. Fortunately, the abovementioned two n-butanol-specific mutants and two isobutanol-specific mutants did not respond to 0–200 mM ethanol (Fig. S4). The apparent *K*_m_ values of mutants W21R/E54V and I183T/D273N towards ethanol were 228 and 174 mM, respectively, which were 23.5- and 17.9-fold higher than that of the wild-type. Similarly, mutants M94V/F272L and S240P possess 13.6- and 12.1-fold higher apparent *K*_m_ towards ethanol when compared with the wild type, respectively. The insensitivity of these mutants to ethanol was maintained at a concentration of 800 mM (Fig. 5E). The complexes in Fig. S5 show that there was no interaction between any of the abovementioned mutants and ethanol. Notably, W21R/E54V and M94V/F272L maintained their respective specificities even with background interference of 500 mM ethanol (Fig. 5F). In addition, we tested the response of special BmoR mutants to other alcohols including methanol, glycol, isopropanol, isopentanol, and 2-methyl-1-pentanol. Fig. S6 shows that mutant T12N could respond to the abovementioned alcohols as well as wild-type BmoR, and mutant W21R/E54V has a narrower signal molecule range when compared with wild-type BmoR, suggesting that mutations in the mutants influenced the range of signal molecules.

The promiscuity of TFBs is a double-edged sword. On the one hand, promiscuity could provide some benefits. For instance, TFBs with promiscuity can expand the signal molecule spectrum. On the other hand, the signal molecule diversity of TFBs could result in an inaccurate response. Many studies have engineered biosensors to realize specific responses ^28, 29^. Endowing a BmoR-based biosensor with n-butanol specificity or isobutanol specificity is required to expand its application. In addition to the native sensing elements, some *de novo* designed biosensors that assemble specific protein functional domains were also developed ^30^. Based on the analysis of BmoR in this work, the specific domains of BmoR might be served as crucial units for establishing novel biosensors to detect other vital compounds.

## Conclusions

The non-ideal properties of wild-type BmoR limit its applications for the screening of single-higher-alcohol industrial-producing strains or the detection of byproducts in the fermentation process. In this study, we engineered a significant region of BmoR and acquired two n-butanol-specific mutants, two isobutanol-specific mutants, and two ultrasensitive mutants. In addition, a mutant with a wider detection range (0–200 mM) was screened out, which could also be reflected in the isobutanol-producing strain. Notably, we observed that the signal-molecule-specific mutants display insensitivity towards ethanol, indicating that these specific mutants could be used to detect the production of byproducts (butanols) in the ethanol fermentation industry. In summary, this work supplied various desirable BmoR mutants that could be employed for constructing biosensors to screen ideal strains or establish dynamic control systems in the field of metabolic engineering.

## Methods

### Strains, media, and materials

*E. coli* XL10-Gold was used for plasmid construction, screening, and construction of the BmoR N-terminal-based random mutagenesis library. LB medium (10 g/L tryptone, 5 g/L yeast extract, and 10 g/L NaCl) was used for strain incubation and library screening. M9 medium (6 g/L Na_2_HPO_4_, 3 g/L KH_2_PO_4_, 1 g/L NH_4_Cl, and 0.5 g/L NaCl, 1 mM MgSO_4_, 0.1 mM CaCl_2_, 10 mg/L VB1, 4 g/L yeast extract, and 40 g/L glucose) was used for the fermentation experiment. Plasmids pSA65, pSA69, pYH1, and pYH10 were obtained from our previous study ^16, 24^. The details of the strains and plasmids used in this study are listed in Table S1.

### Homology modeling and molecular docking of BmoR mutants with signal molecules

AUTODOCK and Chimera 1.14 were used for homology modeling and molecular docking of BmoR with signal molecules (n-butanol, isobutanol, ethanol, or glycol). The tertiary structure of wild-type BmoR, which was modeled in our previous study ^24^, was used as a template to simulate and dock BmoR mutants with signal molecules. All BmoR mutant models were evaluated by PROCHECK, and all models have satisfactory quality, with over 80% of the residues in the most favored region of the calculated z-scores. A grid box (10 × 17 × 8) encompassing the binding pocket of BmoR was set as the search space to explore suitable substrate-binding regions. The interactions between the mutants and signal molecules were analyzed and are shown in the corresponding figures.

### Establishment of the BmoR N-terminal-based mutagenesis library

Error-prone PCR was carried out on the N-terminus of BmoR to generate an N-terminal-based mutagenesis library. The details of the error-prone PCR method are provided in the Supplementary Material. To carry out site-directed mutagenesis of *bmoR*, a pair of primers, including the desired mutations, was synthesized and used to amplify a linearized fragment from pYH1. The PCR products were purified and digested with DpnI and then transferred into *E. coli* XL10-Gold. The primers used in this study are listed in Table S2.

### High-throughput screening of the mutagenesis library

Single colonies harboring plasmid pYH1 containing wild-type or mutated BmoR were pre-inoculated into 5 mL LB with 100 μg/mL ampicillin and then cultured at 37 °C overnight. Next, 50 μL of the seed culture was transferred into 950 μL LB, which was supplemented with appropriate antibiotics and 10 mM n-butanol or isobutanol in 96-deep-well plates. The cultures were then left at 30 °C for 16 h. To measure the apparent *K*_m_ values of the special BmoR mutants towards the butanols or ethanol, the butanol concentrations in 96-deep-well plates ranged between 0–100 mM, and the ethanol concentrations in 96-deep-well plates ranged between 0–800 mM. A microplate reader (BioTek Cytation 3) was used to detect the OD_600_ values and GFP fluorescence.

The excitation and emission wavelengths were set at 470 nm and 510 nm, respectively. The *bmoR* in the colonies that have distinct GFP/OD_600_ values compared to the wild type were sequenced. Apparent *K_m_* values were estimated using Origin8.5 through non-linear regression of the Michaelis-Menten equation. Fully automatic inverted biological microscope (DMi 8) was used to observe green fluorescence. The entire screening process is illustrated in Fig. 1D. The ratio of n-butanol to isobutanol in a mixture of butanol in the culture is described in the Supplementary Material.

### Fermentation verification of BmoR with a wider detection range

Plasmids pYH10 containing wild-type *bmoR* and pYH10-T12N containing *T12N* were individually introduced into strain TW (JCL260 with pSA65, which contained genes *kivd* and *adhA*, and pSA69, which contained genes *alsS*, *ilvC*, and *ilvD*), resulting in strains TW1 and TW2, respectively. For the fermentation experiment, the single colonies were pre-inoculated into 5 mL LB medium with associated antibiotics at 37 ◻ for 12–14 h. Then, 200 μL of culture was inoculated into 20 mL M9 with 0.1 mM IPTG in a 250 mL screw cap conical flask and left at 30 °C in a shaker at 220 rpm. After 36 h, 40 g/L glucose was added to the culture. Samples were taken every 12 h for GFP/OD_600_ measurements and isobutanol detection. GC analysis of the samples was performed as described in our previous study ^31^.

## AUTHOR INFORMATION

### Autor Contributions

Y.-X.H. generated the idea. Y.-X.H., Z.C. and T.W. designed the project. T.W. and Z.C. carried out the experiments. T.W., Z.C., S.G., C.Z. and Y.-X.H. analyzed the data. T.W., Z.C. and Y.-X.H. wrote the manuscript.

### Notes

The authors declare no competing financial interest.

## ACKNOWLEDGMENTS

Research reported in this publication was supported by the National Key R&D Program of China (grant no. 2019YFA0904104), National Natural Science Foundation of China (grant nos.21908007 and 32000059), the Science and Technology Innovation Program of Chinese Academy of Agricultural Sciences (ASTIP TRIC05), and the Fundamental Research Funds for the Central Universities. We thank Biological & Medical Engineering Core Facilities (Beijing Institute of Technology) for providing advanced equipment.

---

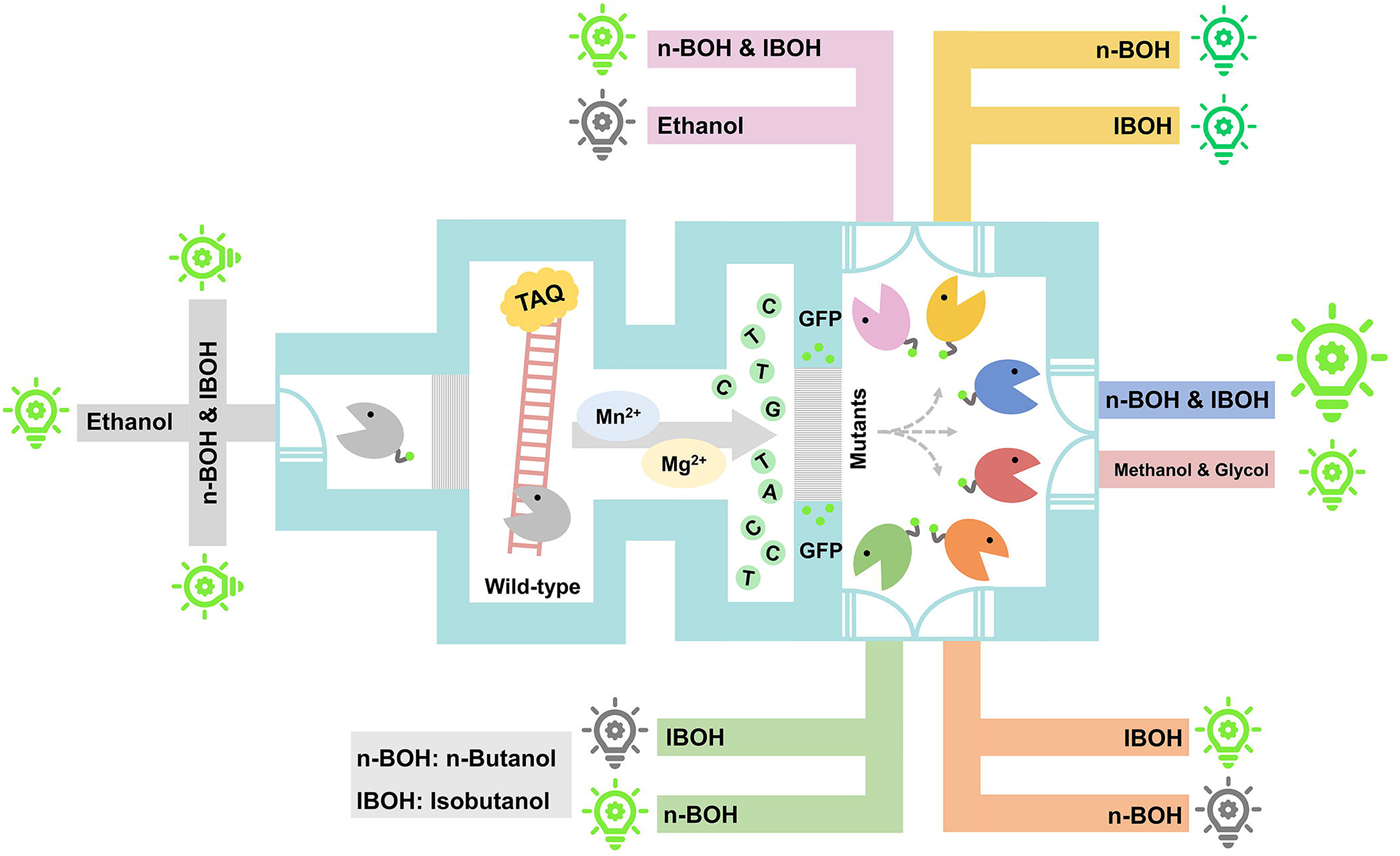

