## Supplementary material for "Engineering transcription factor BmoR mutants for constructing multifunctional alcohol biosensors": Supplmentary

**Supporting Information**

**Contents:**

Materials and methods

Table S1

Table S2

Fig. S1

Fig. S2

Fig. S3

Fig. S4

Fig. S5

Fig. S6

**Materials and methods**

**Establishment of BmoR N-terminal-based mutagenesis library by error-prone PCR**

The first step was to prepare 10× unbalanced dNTPs mixture and this mixture contained 2 mM dATP, 2 mM dGTP, 8 mM dCTP and 8 mM dTTP. The 100 μL reaction mixture contained 1 μL 0.4 ng/L original plasmid pYH1, 10 μL 10× unbalanced dNTPs mixture, 2 μL 25 mM MnCl_2_, 1 μL 200 mM MgCl_2_, 2 μL 10 μM forward primer, 2 μL 10 μM reverse primer, 2 μL 5 U/μL Hieff^TM^ Taq DNA Polymerase, 10 μL 10×M5 Taq PCR Buffer (Mg^2+^ free) and 70 μL ddH_2_O. The amplification program was as follows: 94 °C initial denaturation for 3 min and then 30 cycles of 94 °C denaturation for 1 min, 60 °C annealing for 1 min and 72 °C extension for 2 min. The PCR product was confirmed and purified via agarose gel electrophoresis. The purified *bmoR* N-terminal product was digested by Dpn I at 37 °C for 1-2 h and then cloned into pYH1 by Gibson Assembly. The ligation reaction (total volume 10 μL) containing 3 μL *bmoR* N-terminal fragment after digestion, 2 μL backbone of pYH1 after digestion, and 5 μL Gibson Assembly Mix was left at 50 ℃ for 1 h. Each 5-10 μL of the ligation product was transferred into 50 μL *E. coli* XL10-Gold competent cells.

**Further verification of the response via adding mixed signal molecules**

To verify the response of signal-molecule-specific mutants towards a mixture of signal molecules, the ratio of n-butanol to isobutanol in the culture were set as 1:10 (10 mM : 100 mM),1:8 (10 mM : 80 mM) 1:6 (10 mM : 60 mM), 1:4 (10 mM : 40 mM), 1:2 (10 mM : 20 mM), 1:1 (10 mM : 10 mM), 1:0 (10 mM : 0 mM), 0:1 (0 mM : 10 mM), 1:1 (10 mM : 10 mM), 2:1 (20 mM : 10 mM), 4:1 (40 mM : 10 mM), 6:1 (60 mM : 10 mM), 8:1 (80 mM : 10 mM) or 10:1 (100 mM : 10 mM).

To confirm the insensitivity and specificity of BmoR mutants towards mixed alcohols, the ratio of n-butanol to isobutanol were set as 1:6 (10 mM : 60 mM), 1:4 (10 mM : 40 mM), 1:2 (10 mM : 20 mM), 1:1 (10 mM : 10 mM), 1:0 (10 mM : 0 mM), 0:1 (0 mM : 10 mM), 1:1 (10 mM : 10 mM), 2:1 (20 mM : 10 mM), 4:1 (40 mM : 10 mM) or 6:1 (60 mM : 10 mM) under the noise of 500 mM ethanol in the culture.

**Table S1. Plasmids and strains used in this study**

| **Strain or plasmid** | **Genotype or description** | **Source** |
| --- | --- | --- |
| *E.coli* strains |  |  |
| XL10-Gold | *Tet^r^Δ(mcrA)183 Δ(mcrCB-hsdSMR-mrr)173 endA1 supE44 thi-1 recA1 gyrA96 relA1 lac Hte [F´ proAB lacI^q^ZΔM15 Tn10(Tet^r^) Amy Cam^r^]* | Stratagene |
| JCL260 | *BW25113 rrnBT14 ΔlacZWJ16 hsdR514 ΔaraBADAH33 ΔrhaBADLD78[traD36, proAB+, lacIq ZΔM15, ΔadhE, ΔfrdBC, Δfnr, ΔldhA, Δpta, ΔpflB* | ^1^ |
| TW | JCL260 with pSA65 and pSA69 | This study |
| TW1(wild-type BmoR) | TW with pYH10 | This study |
| TW2(T12N BmoR) | TW with pYH10-T12N | This study |
| Plasmids |  |  |
| pYH1 | *P_bmoR_-bmoR; P_bmo_-gfp; colE1; amp^r^* | ^2^ |
| pYH7 | *P_bmoR_-bmoR; P_bmo_- kan^r^; colE1; amp^r^* | ^2^ |
| pYH10 | *P_bmoR_-bmoR; P_bmo_- kan^r^; colA; cm^r^* | ^2^ |
| pSA65 | *P_L_lacO_1_-kivd-adhA, colE1, amp^r^* | ^3^ |
| pSA69 | *P_L_lacO_1_-alsS-ilvC-ilvD, p15A, kan^r^* | ^3^ |
| pYH10-T12N | pYH10 expressing T12N BmoR | This study |

(1)	Atsumi, S.,Cann, A. F.,Connor, M. R.,Shen, C. R.,Smith, K. M.,Brynildsen, M. P.,Chou, K. J.,Hanai, T.,Liao, J. C., (2008) Metabolic engineering of *Escherichia coli* for 1-butanol production. *Metab. Eng. 10* (6), 305-311.

(2)	Yu, H.,Chen, Z.,Wang, N.,Yu, S.,Yan, Y.,Huo, Y.-X., (2019) Engineering transcription factor BmoR for screening butanol overproducers. *Metab. Eng. 56*, 28-38.

(3)	Atsumi, S.,Hanai, T.,Liao, J. C., (2008) Non-fermentative pathways for synthesis of branched-chain higher alcohols as biofuels. *Nature 451* (7174), 86-89.

**Table S2.** Primers used in this study

| Primer | Sequence (5’-3’) | Usage |
| --- | --- | --- |
| WT-001 | gtcgagaatcggtctgatccg | For introducing  random mutations into  N-terminal of *bmoR* |
| WT-002 | cggtgaaagcaccttcttcg |  |
| WT-091 | aattcgctcgtctggaaaacgttgcttctatgcgtc | For introducing the mutation of T12N into pYH10 |
| WT-092 | aagcaacgttttccagacgagcgaattcctgc |  |
| R211K-R | aggtctgTTTgaacagctggttttcgatga | For site-saturation mutagenesis at Arg211 of *bmoR* |
| R211C-R | aggtctgTGCgaacagctggttttcgat |  |
| R211D-R | aggtctgGACgaacagctggttttcgat |  |
| R211E-R | aggtctgGAAgaacagctggttttcgat |  |
| R211F-R | aggtctgTTCgaacagctggttttcgat |  |
| R211H-R | aggtctgCACgaacagctggttttcgat |  |
| R211I-R | aggtctgATAgaacagctggttttcgat |  |
| R211L-R | aggtctgCTAgaacagctggttttcgat |  |
| R211N-R | aggtctgAACgaacagctggttttcgat |  |
| R211P-R | aggtctgCCAgaacagctggttttcgat |  |
| R211Q-R | aggtctgCAAgaacagctggttttcgat |  |
| R211S-R | aggtctgAGCgaacagctggttttcgat |  |
| R211T-R | aggtctgACAgaacagctggttttcgat |  |
| R211V-R | aggtctgGTAgaacagctggttttcgat |  |
| R211W-R | aggtctgTGGgaacagctggttttcgat |  |
| R211Y-R | aggtctgTACgaacagctggttttcgat |  |
| R211G-R | aggtctgGGAgaacagctggttttcga |  |
| R211M-R | aggtctgATGgaacagctggttttc |  |
| R211K-F | ccagctgttcAAAcagaccttcgctgactg |  |
| R211C-F | ccagctgttcTGCcagaccttcgctgact |  |
| R211D-F | ccagctgttcGACcagaccttcgctgact |  |
| R211E-F | ccagctgttcGAAcagaccttcgctgact |  |
| R211F-F | ccagctgttcTTCcagaccttcgctgact | For site-saturation mutagenesis at Arg211 of *bmoR* |
| R211G-F | ccagctgttcGGAcagaccttcgctgACT |  |
| R211H-F | ccagctgttcCACcagaccttcgctgact |  |
| R211I-F | ccagctgttcATAcagaccttcgctgact |  |
| R211L-F | ccagctgttcCTAcagaccttcgctgact |  |
| R211M-F | ccagctgttcATGcagaccttcgctgact |  |
| R211N-F | ccagctgttcAACcagaccttcgctgact |  |
| R211P-F | ccagctgttcCCAcagaccttcgctgact |  |
| R211Q-F | ccagctgttcCAAcagaccttcgctgact |  |
| R211S-F | ccagctgttcAGCcagaccttcgctgact |  |
| R211T-F | ccagctgttcACAcagaccttcgctgact |  |
| R211V-F | ccagctgttcGTAcagaccttcgctgact |  |
| R211Y-F | ccagctgttcTACcagaccttcgctgact |  |
| R211W-F | ccagctgttcTGGcagaccttcgct |  |
| Q212G-F | gctgttccgtGGTaccttcgctgact | For site-saturation mutagenesis at Gln212 of *bmoR* |
| Q212I-F | gctgttccgtATTaccttcgctgact |  |
| Q212L-F | gctgttccgtCTTaccttcgctgact |  |
| Q212P-F | gctgttccgtCCTaccttcgctgact |  |
| Q212V-F | gctgttccgtGTTaccttcgctgact |  |
| Q212F-F | gctgttccgtTTTaccttcgctgact |  |
| Q212W-F | gctgttccgtTGGaccttcgctgact |  |
| Q212Y-F | gctgttccgtTATaccttcgctgact |  |
| Q212D-F | gctgttccgtGATaccttcgctgact |  |
| Q212E-F | gctgttccgtGAAaccttcgctgact |  |
| Q212R-F | gctgttccgtCGTaccttcgctgact |  |
| Q212H-F | gctgttccgtCATaccttcgctgact |  |
| Q212K-F | gctgttccgtAAAaccttcgctgact |  |
| Q212S-F | gctgttccgtTCTaccttcgctgact |  |
| Q212T-F | gctgttccgtACTaccttcgctgact |  |
| Q212C-F | gctgttccgtTGTaccttcgctgact | For site-saturation mutagenesis at Gln212 of *bmoR* |
| Q212M-F | gctgttccgtATGaccttcgctgact |  |
| Q212N-F | gctgttccgtAATaccttcgctgact |  |
| Q212N-R | gcgaaggtAATacggaacagctggttttc |  |
| Q212M-R | gcgaaggtATGacggaacagctggttttc |  |
| Q212C-R | gcgaaggtTGTacggaacagctggttttc |  |
| Q212T-R | gcgaaggtACTacggaacagctggttttc |  |
| Q212S-R | gcgaaggtTCTacggaacagctggttttc |  |
| Q212K-R | gcgaaggtAAAacggaacagctggttttc |  |
| Q212H-R | gcgaaggtCATacggaacagctggttttc |  |
| Q212R-R | gcgaaggtCGTacggaacagctggttttc |  |
| Q212E-R | gcgaaggtGAAacggaacagctggttttc |  |
| Q212D-R | gcgaaggtGATacggaacagctggttttc |  |
| Q212Y-R | gcgaaggtTATacggaacagctggttttc |  |
| Q212W-R | gcgaaggtTGGacggaacagctggttttc |  |
| Q212F-R | gcgaaggtTTTacggaacagctggttttc |  |
| Q212V-R | gcgaaggtGTTacggaacagctggttttc |  |
| Q212P-R | gcgaaggtCCTacggaacagctggttttc |  |
| Q212L-R | gcgaaggtCTTacggaacagctggttttc |  |
| Q212I-R | gcgaaggtATTacggaacagctggttttc |  |
| Q212G-R | gcgaaggtGGTacggaacagctggttttc |  |
| N259E-F | aatcgctggtctgGAActggaagctgttgct | For site-saturation mutagenesis at Asn259 of *bmoR* |
| N259G-F | aatcgctggtctgGGcctggaagctgttgc |  |
| N259I-F | aatcgctggtctgaTcctggaagctgttgc |  |
| N259L-F | aatcgctggtctgCTcctggaagctgttgc |  |
| N259P-F | aatcgctggtctgCCcctggaagctgttgc |  |
| N259V-F | aatcgctggtctgGTcctggaagctgttgc |  |
| N259F-F | aatcgctggtctgTTcctggaagctgttgc |  |
| N259Y-F | aatcgctggtctgTACctggaagctgttgc |  |
| N259D-F | aatcgctggtctgGacctggaagctgttgc | For site-saturation mutagenesis at Asn259 of *bmoR* |
| N259R-F | aatcgctggtctgCGCctggaagctgttgc |  |
| N259H-F | aatcgctggtctgCAcctggaagctgttgc |  |
| N259K-F | aatcgctggtctgaaActggaagctgttgc |  |
| N259S-F | aatcgctggtctgTCcctggaagctgttgc |  |
| N259T-F | aatcgctggtctgaCcctggaagctgttgc |  |
| N259C-F | aatcgctggtctgTGcctggaagctgttgc |  |
| N259M-F | aatcgctggtctgaTGctggaagctgttgc |  |
| N259Q-F | aaaatcgctggtctgCAActggaagctgttg |  |
| N259W-F | ctgaaaatcgctggtctgTGGctggaagctgttgct |  |
| N259W-R | CCAcagaccagcgattttcagaccagcacggttca |  |
| N259M-R | ccagCAtcagaccagcgattttcagaccagcac |  |
| N259C-R | ccaggCAcagaccagcgattttcagaccagcac |  |
| N259T-R | ccaggGtcagaccagcgattttcagaccagcac |  |
| N259S-R | ccaggGAcagaccagcgattttcagaccagcac |  |
| N259K-R | ccagTttcagaccagcgattttcagaccagcac |  |
| N259H-R | ccaggtGcagaccagcgattttcagaccagcac |  |
| N259R-R | ccaggCGcagaccagcgattttcagaccagcac |  |
| N259D-R | ccaggtAcagaccagcgattttcagaccagcac |  |
| N259Y-R | ccaggtAcagaccagcgattttcagaccagcac |  |
| N259F-R | ccaggAAcagaccagcgattttcagaccagcac |  |
| N259V-R | ccaggACcagaccagcgattttcagaccagcac |  |
| N259P-R | ccaggGGcagaccagcgattttcagaccagcac |  |
| N259L-R | ccaggAGcagaccagcgattttcagaccagcac |  |
| N259I-R | ccaggAtcagaccagcgattttcagaccagcac |  |
| N259G-R | ccaggCCcagaccagcgattttcagaccagcac |  |
| N259Q-R | ttccagTTGcagaccagcgattttcagaccagcac |  |
| N259E-R | cttccagTTCcagaccagcgattttcagaccagcac |  |
| E261P-F | tggtctgaacctgCCTgctgttgctgaccaccgtt | For site-saturation mutagenesis at Glu261 of *bmoR* |
| E261A-F | tggtctgaacctggCTgctgttgctgaccacc |  |
| E261G-F | tggtctgaacctggGTgctgttgctgaccacc |  |
| E261I-F | gctggtctgaacctgATTgctgttgctgaccacc |  |
| E261L-F | gctggtctgaacctgCTTgctgttgctgaccacc |  |
| E261V-F | gctggtctgaacctggTTgctgttgctgaccacc |  |
| E261F-F | gctggtctgaacctgTTTgctgttgctgaccacc |  |
| E261W-F | gctggtctgaacctgTGGgctgttgctgaccacc |  |
| E261Y-F | gctggtctgaacctgTATgctgttgctgaccacc |  |
| E261D-F | gctggtctgaacctgGATgctgttgctgaccacc |  |
| E261R-F | gctggtctgaacctgCGTgctgttgctgaccacc |  |
| E261H-F | gctggtctgaacctgCATgctgttgctgaccacc |  |
| E261K-F | gctggtctgaacctgAAAgctgttgctgaccacc |  |
| E261S-F | gctggtctgaacctgTCTgctgttgctgaccacc |  |
| E261T-F | gctggtctgaacctgACTgctgttgctgaccacc |  |
| E261C-F | gctggtctgaacctgTGTgctgttgctgaccacc |  |
| E261M-F | gctggtctgaacctgATGgctgttgctgaccacc |  |
| E261N-F | gctggtctgaacctgAATgctgttgctgaccacc |  |
| E261Q-F | gctggtctgaacctgCaagctgttgctgaccacc |  |
| E216Q-R | gcaacagcttGcaggttcagaccagcgattttcagacc |  |
| E216N-R | gcaacagcATTcaggttcagaccagcgattttcagacc |  |
| E216M-R | gcaacagcCATcaggttcagaccagcgattttcagacc |  |
| E216C-R | gcaacagcACAcaggttcagaccagcgattttcagacc |  |
| E216T-R | gcaacagcAGTcaggttcagaccagcgattttcagacc |  |
| E216S-R | gcaacagcAGAcaggttcagaccagcgattttcagacc |  |
| E216K-R | gcaacagcTTTcaggttcagaccagcgattttcagacc |  |
| E216R-R | gcaacagcACGcaggttcagaccagcgattttcagacc |  |
| E216F-R | gcaacagcAAAcaggttcagaccagcgattttcagacc |  |
| E216G-R | cagcACccaggttcagaccagcgattttcagac |  |
| E216A-R | cagcAGccaggttcagaccagcgattttcagac | For site-saturation mutagenesis at Glu261 of *bmoR* |
| E216V-R | aacagcAACcaggttcagaccagcgattttcagac |  |
| E216H-R | gcaacagcATGcaggttcagaccagcgattttcagac |  |
| E216D-R | gcaacagcATCcaggttcagaccagcgattttcagac |  |
| E216Y-R | gcaacagcATAcaggttcagaccagcgattttcagac |  |
| E216W-R | gcaacagcCCAcaggttcagaccagcgattttcagac |  |
| E216P-R | gcaacagcAGGcaggttcagaccagcgattttcagac |  |
| E216L-R | gcaacagcAAGcaggttcagaccagcgattttcagac |  |
| E216I-R | gcaacagcAATcaggttcagaccagcgattttcagac |  |
| E261P-F | tggtctgaacctgCCTgctgttgctgaccaccgtt |  |
| E261A-F | tggtctgaacctggCTgctgttgctgaccacc |  |
| E261G-F | tggtctgaacctggGTgctgttgctgaccacc |  |
| E261I-F | gctggtctgaacctgATTgctgttgctgaccacc |  |
| E261L-F | gctggtctgaacctgCTTgctgttgctgaccacc |  |
| E261V-F | gctggtctgaacctggTTgctgttgctgaccacc |  |

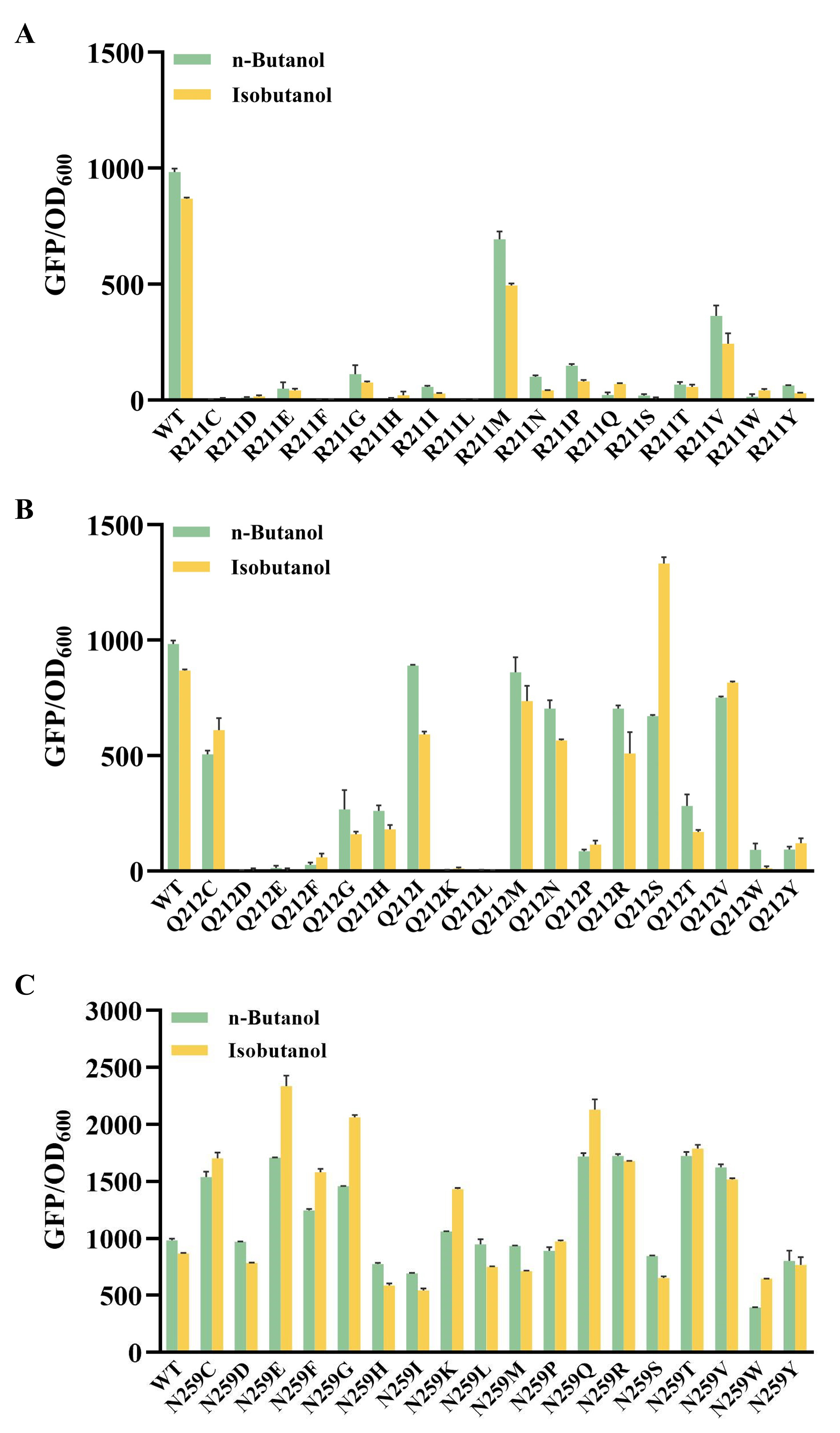

**Fig. S1.** Site-saturation mutagenesis of *bmoR*. **(A)** Site-saturation mutagenesis at Arg211 of *bmoR*. **(B)** Site-saturation mutagenesis at Gln212 of *bmoR*. **(C)** Site-saturation mutagenesis at Asn259 of *bmoR*.

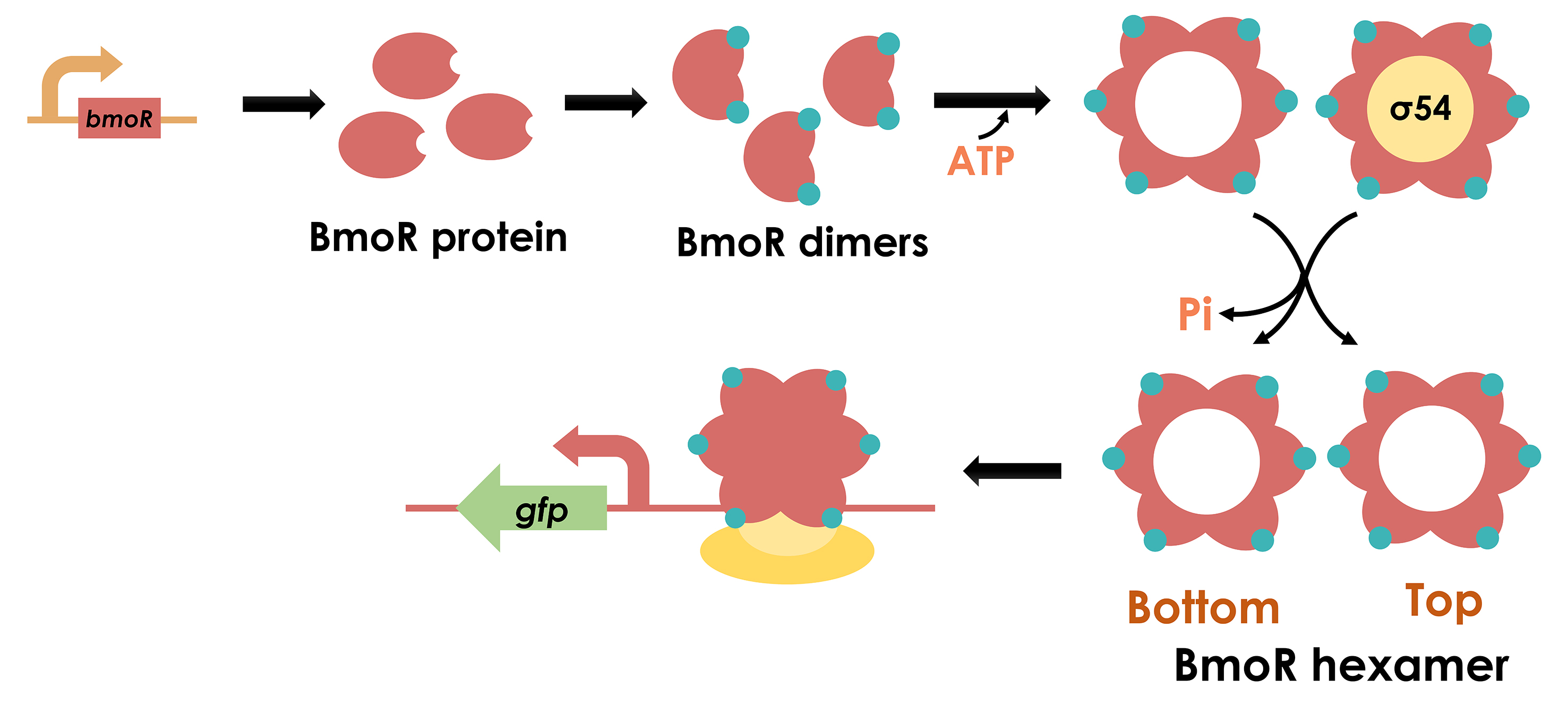
**Fig. S2.** The whole activation process of BmoR.

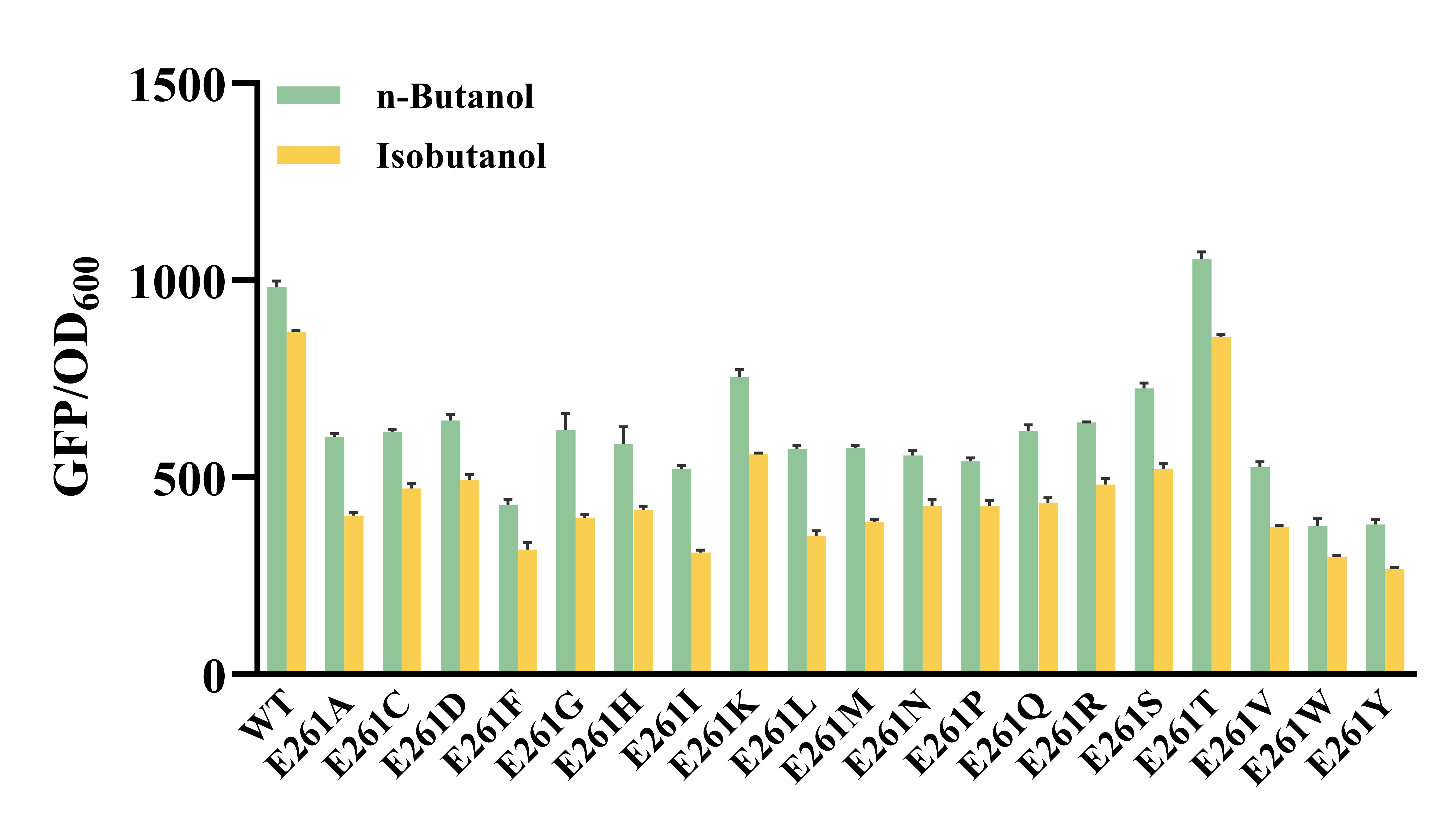
**Fig. S3.** Site-saturation mutagenesis at Glu261 of mutant W21R/E54V.

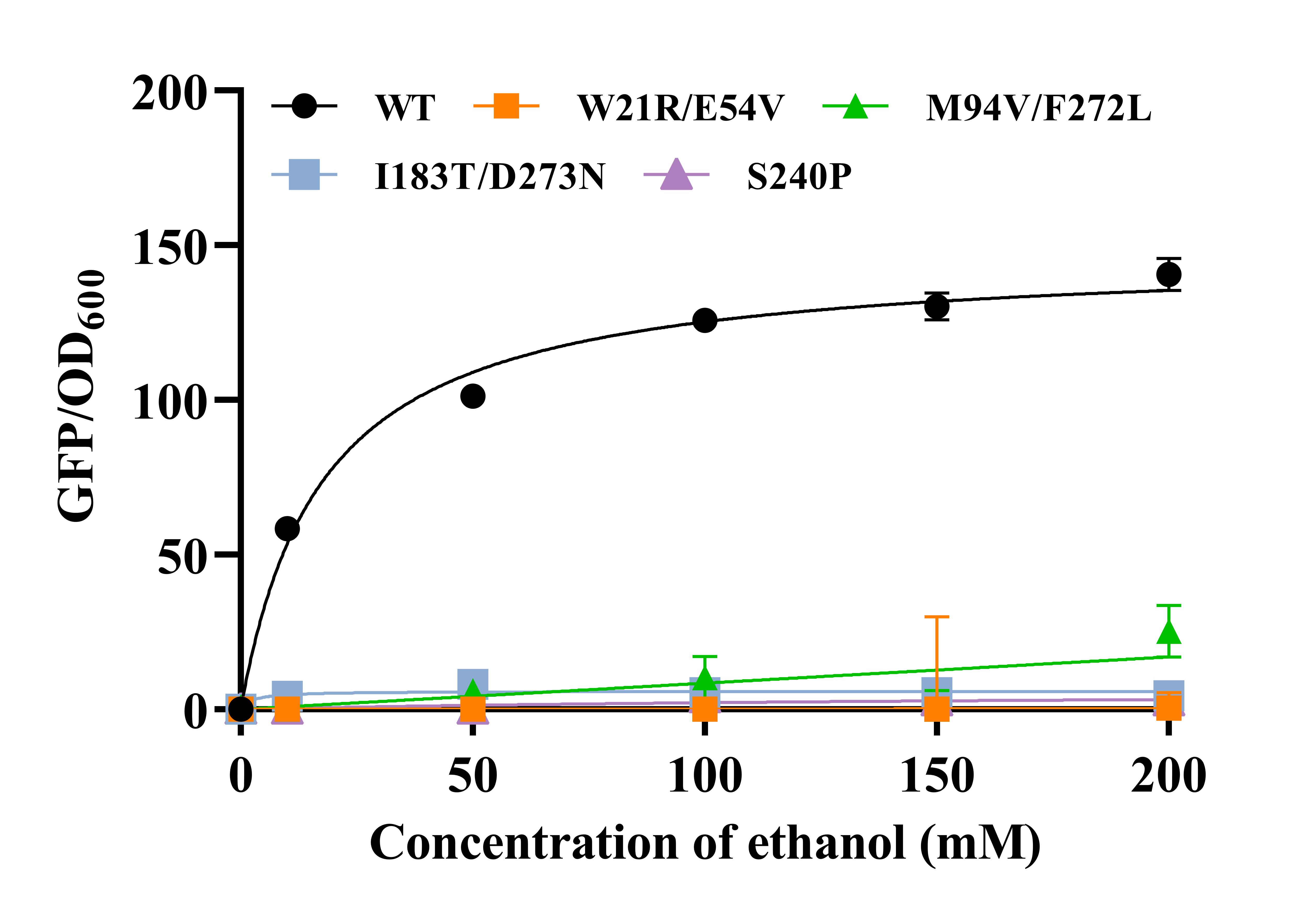

**Fig. S4.** The response values of wild-type BmoR and n-butanol-specific mutants(W21R/E54V and I183T/D273N) and isobutanol-specific mutants (M94V/F272L and S240P) to 0-200 mM ethanol.

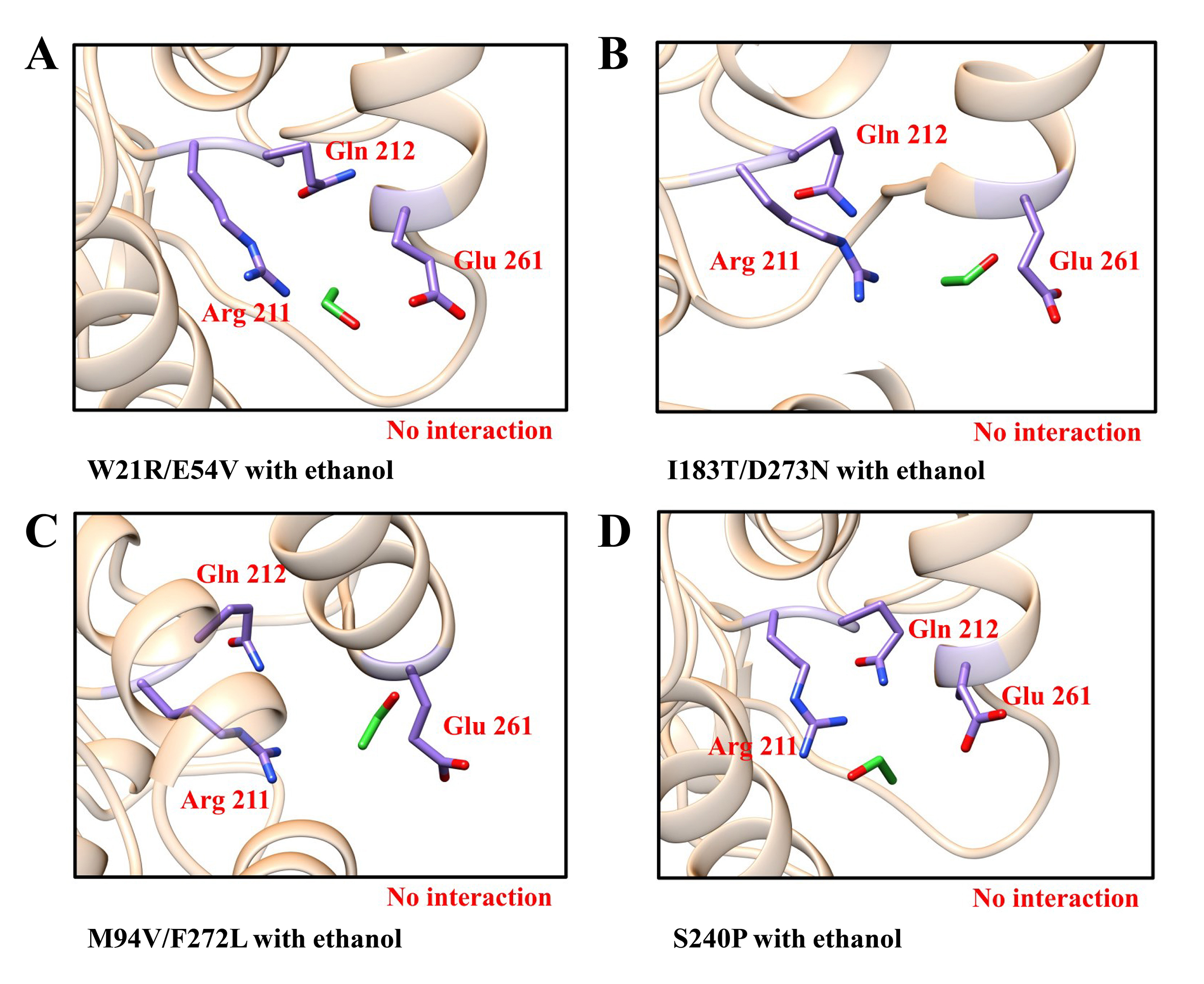
**Fig. S5.** Molecular docking of BmoR mutants with ethanol. **(A)** The complex of W21R/E54V with ethanol. **(B)** The complex of I183T/D273N with ethanol. **(C)** The complex of M94V/F272L with ethanol. **(D)** The complex of S240P with ethanol.

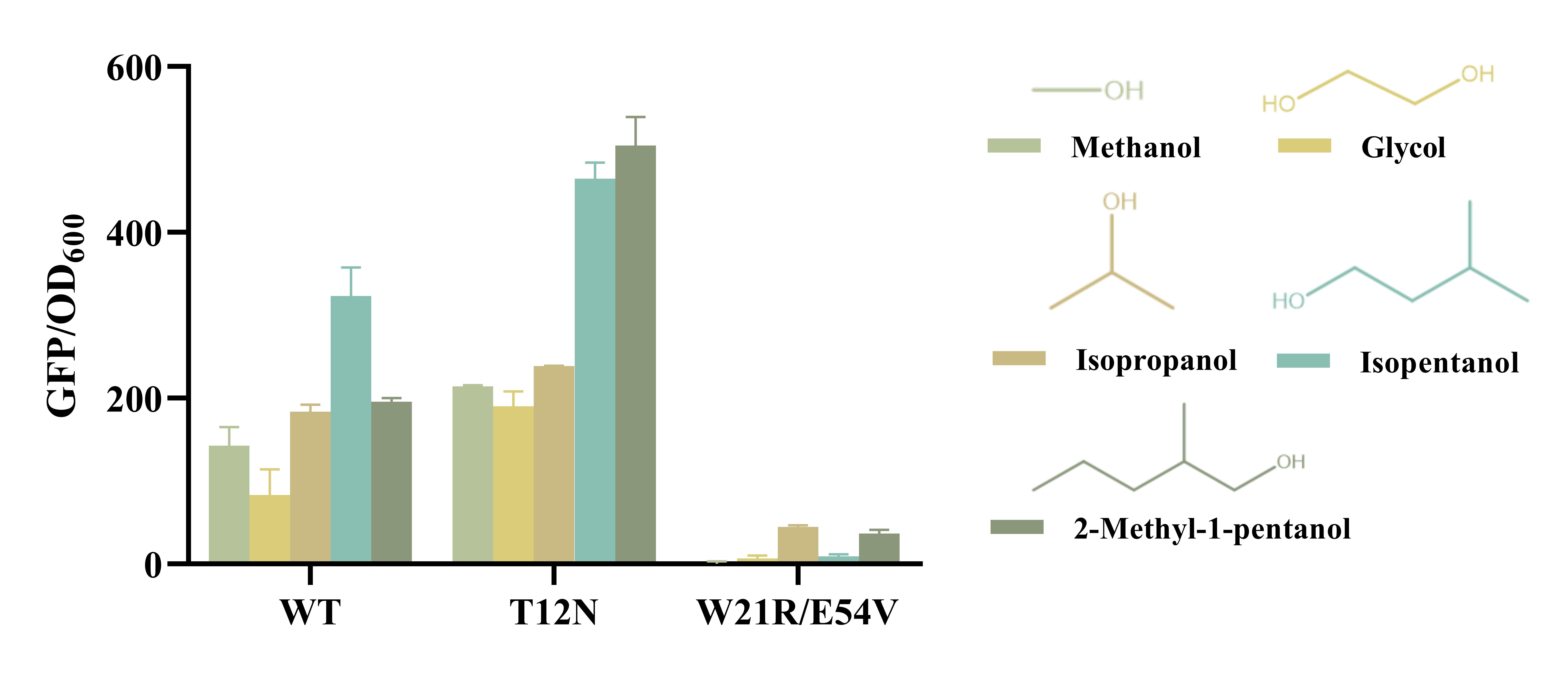

**Fig. S6.** The response values of wild-type BmoR and mutants (T12N and W21R/E54V BmoR) towards other alcohols including methanol, glycol, isopropanol, isopentanol and 2-methyl-1-pentanol.
